# Enhanced C/EBPα function extends healthspan and lifespan in the African turquoise killifish

**DOI:** 10.1101/2025.02.18.638802

**Authors:** Christine Müller, Joscha S. Muck, Gertrud Kortman, Josephine Hartung, Eugene Berezikov, Cornelis F. Calkhoven

## Abstract

The transcription factor CCAAT/enhancer binding protein alpha (C/EBPα) regulates cell differentiation, proliferation, and function in various tissues, including the liver, adipose tissue, skin, lung, and hematopoietic system. Studies in rats, mice, humans, and chickens have shown that CEBPA mRNA undergoes alternative translation initiation, producing three C/EBPα protein isoforms. Two of these isoforms act as full-length transcription factors with N-terminal transactivation domains and a C-terminal dimerization and DNA-binding domains. The third isoform is an N-terminally truncated variant, translated from a downstream AUG codon. It competes with full-length isoforms for DNA binding, thereby antagonizing their activity. Expression of the truncated C/EBPα isoform depends on the initial translation of a short upstream open reading frame (uORF) in CEBPA mRNA and subsequent re-initiation at a downstream AUG codon, a process stimulated by mTORC1 signaling. We investigated whether the ortholog of the CEBPA gene in the evolutionarily distant, short-lived African turquoise killifish (*Nothobranchius furzeri*) is regulated by similar mechanisms. Our findings reveal that the uORF-mediated regulation of C/EBPα isoform expression is conserved in killifish. Disruption of the uORF selectively eliminates the truncated isoform, leading to unrestrained activity of the full-length C/EBPα isoforms. This genetic modification significantly extended both the median and maximal lifespan and improved the healthspan of male *N. furzeri*. These results highlight a conserved mechanism of *CEBPA* gene regulation across species and its potential role in modulating the lifespan and aging phenotypes.

## INTRODUCTION

The African turquoise killifish, *Nothobranchius furzeri,* is a unique vertebrate species that has attracted considerable attention in the field of aging research because of its exceptionally short lifespan. Native to ephemeral ponds in the semi-arid regions of Mozambique and Zimbabwe, this species has evolved to complete its life cycle rapidly, often within a few months, to adapt to the transient availability of water in its natural habitat. These killifish show a pronounced and well-characterized aging process in captivity, displaying many of the hallmarks of aging observed in longer-lived vertebrates, including declines in cognitive and physical performance, changes in tissue integrity, increase in cancer incidence, and alterations in gene expression profiles (Genade et al., 2005; Hu & Brunet, 2018; Reichard, Cellerino, & Valenzano, 2015). *N. furzeri* has a fully sequenced genome, and genetic tools have been developed, including CRISPR/Cas9-mediated gene editing (Bedbrook, Nath, Nagvekar, Deisseroth, & Brunet, 2023; Cui, Willemsen, & Valenzano, 2020; Harel, Valenzano, & Brunet, 2016; Petzold et al., 2013; Reichwald et al., 2015)(see for genome browser https://nfingb.leibniz-fli.de).

The CCAAT/enhancer binding protein family of basic region leucine-zipper (bZIP) transcription factors consists of six members (designated α to ζ) that are widely expressed and involved in cell proliferation, differentiation, metabolism, and senescence (Lopes-Paciencia et al., 2019; Nerlov, 2007; Ramji & Foka, 2002). C/EBPs regulate gene transcription by forming dimers and binding to C/EBP- specific recognition sites in the genome. The expression of C/EBPα and C/EBPβ proteins is uniquely regulated at the level of mRNA translation, involving a *cis*-regulatory upstream open reading frame (uORF) and translation into three protein isoforms of different lengths from a single mRNA molecule (Calkhoven, Bouwman, Snippe, & Ab, 1994; Calkhoven, Muller, & Leutz, 2000). From long to short, C/EBPα comprises extended-, full-length(p42)-, and truncated(p30)-C/EBPα isoforms. The extended-C/EBPα is translated from an unconventional CUG codon in most vertebrates, or a GUG codon in humans. Its expression is generally low, and it plays a more specialized role in activating Polymerase I-controlled ribosomal DNA transcription and the regulation of cell size (Muller, Bremer, Schreiber, Eichwald, & Calkhoven, 2010). The p42 isoform is translated from the first AUG codon in the *CEBPA* reading frame and functions as a full transactivating factor. It contains an N-terminal transactivation domain and protein-protein interaction domains, as well as a C-terminal dimerization and DNA-binding domain. P42-C/EBPα is involved in activating Polymerase II-mediated transcription of specific target genes. The third p30-C/EBPα isoform is translated from a downstream in-frame AUG codon, which depends on the preceding translation of the uORF and the subsequent re-initiation at the downstream AUG codon. This isoform lacks most of the N-terminal domains for transactivation but retains the C-terminal dimerization and DNA-binding domains. By competing with p42-C/EBPα for DNA binding, it acts as a competitive inhibitor for p42-C/EBPα. Mutation of the uORF abolishes p30-C/EBPα expression, relieving its inhibitory action on p42-C/EBPα and resulting in the unrestrained activity of p42-C/EBPα (Calkhoven et al., 1994; Ossipow, Descombes, & Schibler, 1993).

The present study was preceded by investigations of the regulation and function of C/EBPβ isoforms (Calkhoven et al., 2000; Zidek et al., 2015). These studies showed that mutation of the uORF (ΔuORF) in the *CEBPB* gene in cell lines or mice similarly abolishes the expression of the truncated C/EBPβ isoform called C/EBPβ-LIP, thereby unleashing the transactivating function of the full-length C/EBPβ-LAP isoform, resulting in C/EBPβ superfunction. C/EBPβ-ΔuORF mutation in mice induces calorie-restriction type metabolic phenotypes, reduces spontaneous cancer incidence, and extends the median (20%) and maximum (10%) lifespan, the latter only in females (Muller et al., 2018; Muller et al., 2022; Zidek et al., 2015).

Here, we investigated whether different *N. furzeri* (Nf)C/EBPα isoforms are expressed in killifish and whether the truncated NfC/EBPα isoform depends on uORF translation, as observed in higher vertebrates. Our findings demonstrate that uORF-mediated regulation of NfC/EBPα isoforms is conserved in *N. furzeri*. Disruption of uORF function selectively abolishes the truncated NfC/EBPα isoform, leading to enhanced transcriptional activity of the full-length isoform. *In vivo*, this genetic modification significantly extends both median and maximal lifespan while improving certain healthspan parameters in males. These benefits were absent in females, indicating a sex-specific response to uORF-dependent regulation of NfC/EBPα proteins in *N. furzeri*.

## RESULTS

### CEBPA mRNA and C/EBPα protein structures are conserved in vertebrates

The conservation of the mRNA primary structure throughout vertebrate evolution is remarkable, as demonstrated by the alignment of mRNA sequences of killifish with those from human, mouse, chicken, Xenopus, and the ancient coelacanth (**Figure 1A-D**). It encompasses the precise position of the uORF, distribution of translation initiation sites, and their relative strength, as determined by their adherence to the Kozak consensus sequence (Kozak, 1984). This conservation predicts the expression of both extended and full-length isoforms as well as the uORF-dependent translation of a truncated isoform through mechanisms similar to those described for orthologous transcripts in other vertebrates (Calkhoven et al., 2000).

**Figure 1.**
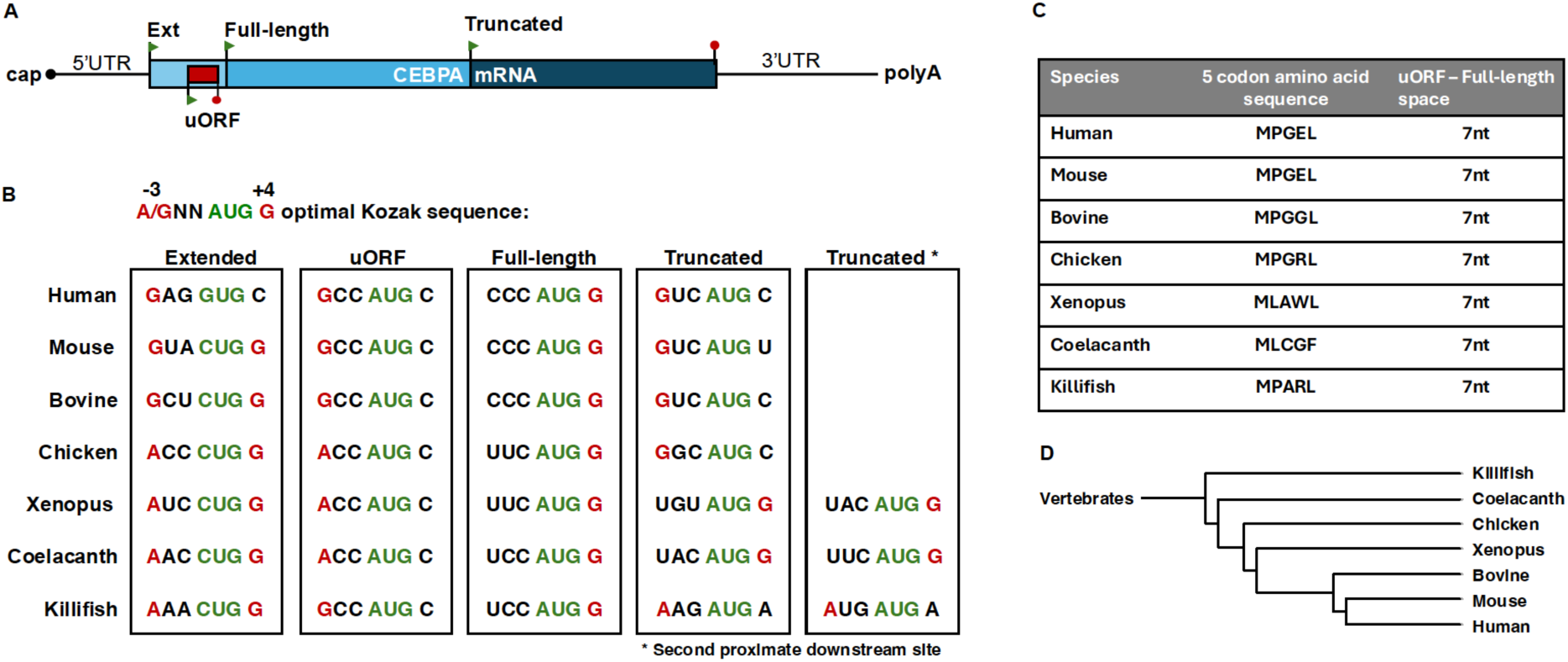
Evolutionary conservation of the CEBPA mRNA primary structure. (**A**) Schematic representation of CEBPA mRNA with 5’cap, 5’ and 3’ untranslated regions (UTR), translation initiation sites (green arrow points) for extended (Ext), full-length and truncated C/EBPα protein isoforms, the upstream open reading frame (uORF), and polyadenylation tail (polyA). The uORF is +2nt out of frame with the CEBPA coding frame. (**B**) Comparison of CEBPA translation initiation sites in different species and their adherence to the Kozak consensus sequence for optimal translation initiation shown above. (**C**) Pentapeptide sequences of uORFs from different species and the space between the stop codon uORF and the initiation codon for full-length C/EBPα in nucleotides (nt). (**D**) Phylogenetic tree of the depicted species (generated with https://phylot.biobyte.de)

Alignment of the NfC/EBPα protein sequence, derived from the genomic transcript of the *NfCEBPA* gene, with C/EBPα protein sequences from human, mouse, and chicken revealed strong conservation in the carboxy-terminal region. This region contains a basic DNA-binding domain and a leucine zipper dimerization domain (bZIP domain) (**Figure 2**) (Gene ID: 107381504; (Petzold et al., 2013)). The amino-terminal region also contains conserved sequences associated with transactivation, specifically the domain spanning amino acids 36 and 73 in the killifish sequence. This domain is unique to C/EBPα and absent in other C/EBP family members, confirming the identity of the sequence as C/EBPα. Two methionine residues, located downstream of the transactivation domain, are predicted to serve as initiation sites for the truncated isoform. Additionally, an upstream CUG codon in a favorable Kozak context is predicted to function as an initiation site for an N-terminally extended isoform, with similar initiation codons observed across species (**Figure 1B**).

**Figure 2.**
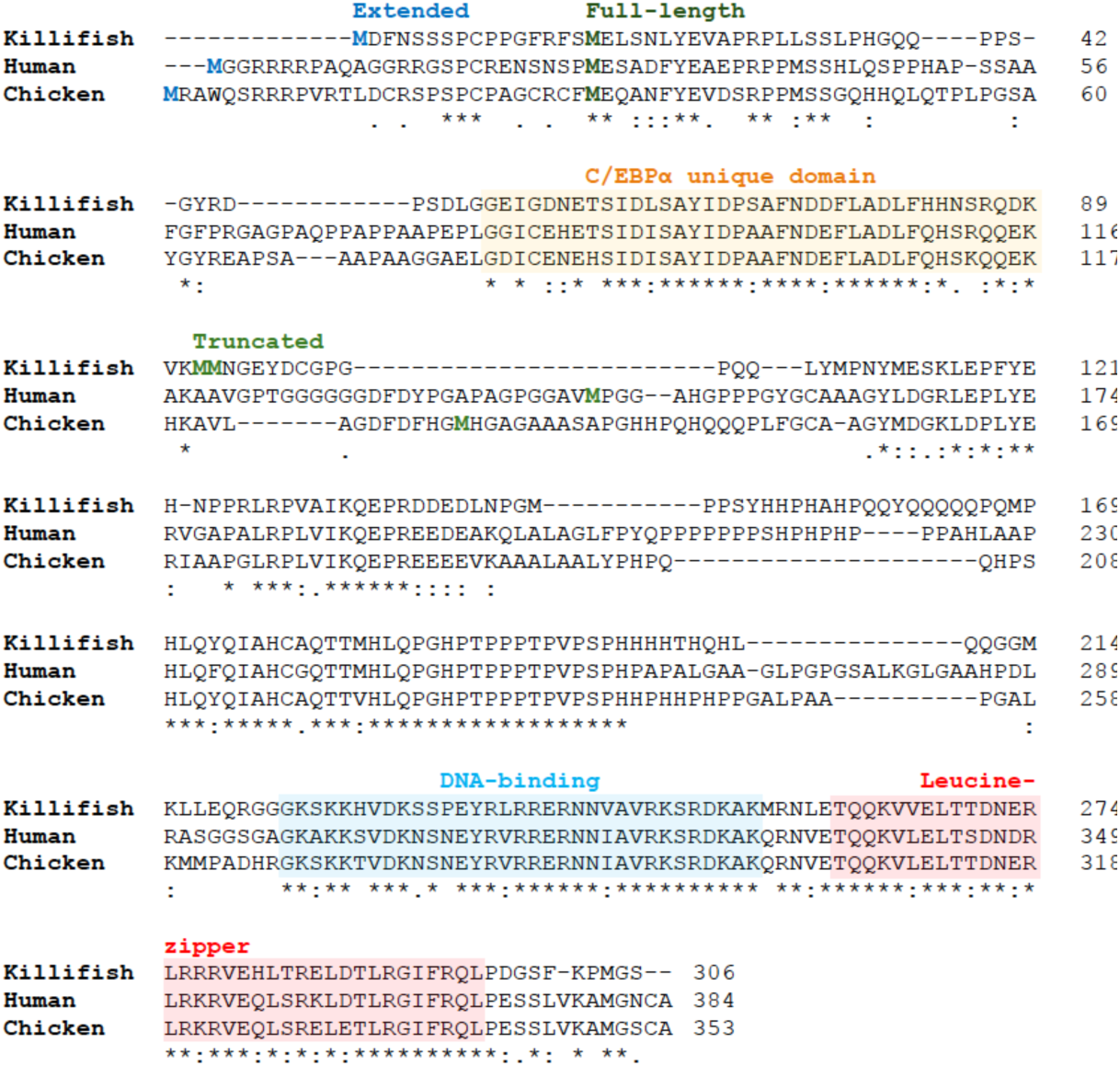
C/EBPα sequence alignment. Amino acids are aligned and marked conserved (*), conservative changes (:), and semiconservative changes (.). The initiating methionines for the Extended isoform are highlighted in blue, corresponding to the alternative start codons CUG (in killifish and chicken) or GUG (in humans). The initiating methionines for Full-length and Truncated isoforms are highlighted in dark and light green, respectively. The unique N-terminal transactivation domain of C/EBPα is marked yellow, and the C-terminal conserved DNA-binding and leucine-zipper dimerization domains (bZIP) are marked in blue and red, respectively.

### uORF-dependent translation of the NfCEBPA mRNA into three protein isoforms

To analyze NfC/EBPα protein isoform expression, we cloned the *NfCEBPA* cDNA sequence into the eukaryotic expression vectors pCDNA3.1 and pSG5. Owing to the lack of anti-NfC/EBPα antibodies, a hemagglutinin (HA) epitope was added at the C-terminus. Transfection of the wild-type (wt) cDNA in COS-1, HEK293, and Hepa1-6 cell lines resulted in the expression of the three expected protein isoforms, labeled extended (Ext), full-length (Fl), and truncated (Tr) (**Figure 3A)**. Next, we examined the usage of the predicted translation initiation sites and assessed the consequences of their perturbation on the expression of NfC/EBPα isoform in COS-1 cells (**Figure 3B**). Removal of the entire 5’UTR (Δ5’UTR) led to the exclusive expression of Fl-NfC/EBPα, indicating that the regulatory sequences required for additional isoform expression reside within the 5’UTR. Mutation of uORF-ATG into ATC (ΔATG) resulted in the loss of Tr-NfC/EBPα expression, while converting uORF-ATG into an optimal Kozak sequence (GCC ATG C-> GCC ATG G) (ATG^Opt^) caused upregulation of Tr-NfC/EBPα, confirming that its expression is dependent on uORF translation. Mutation of the Ext-CTG codon into CTC (ΔCTG) resulted in the loss of Ext-NfC/EBPα, proving its use as an alternative initiation codon. Finally, mutation of the double ATG (ΔATG^Tr^), believed to be the initiation sites for the Tr-NfC/EBPα isoform, caused a shift in expression to a smaller protein, likely due to ribosomes scanning to the next available AUG codon in the *NfCEBPA* reading frame. This confirmed that the double ATGs serve as initiation sites for Tr-NfC/EBPα expression. The expression of single Tr-NfC/EBPα proteins is shown for reference (**Figure 3B**).

**Figure 3.**
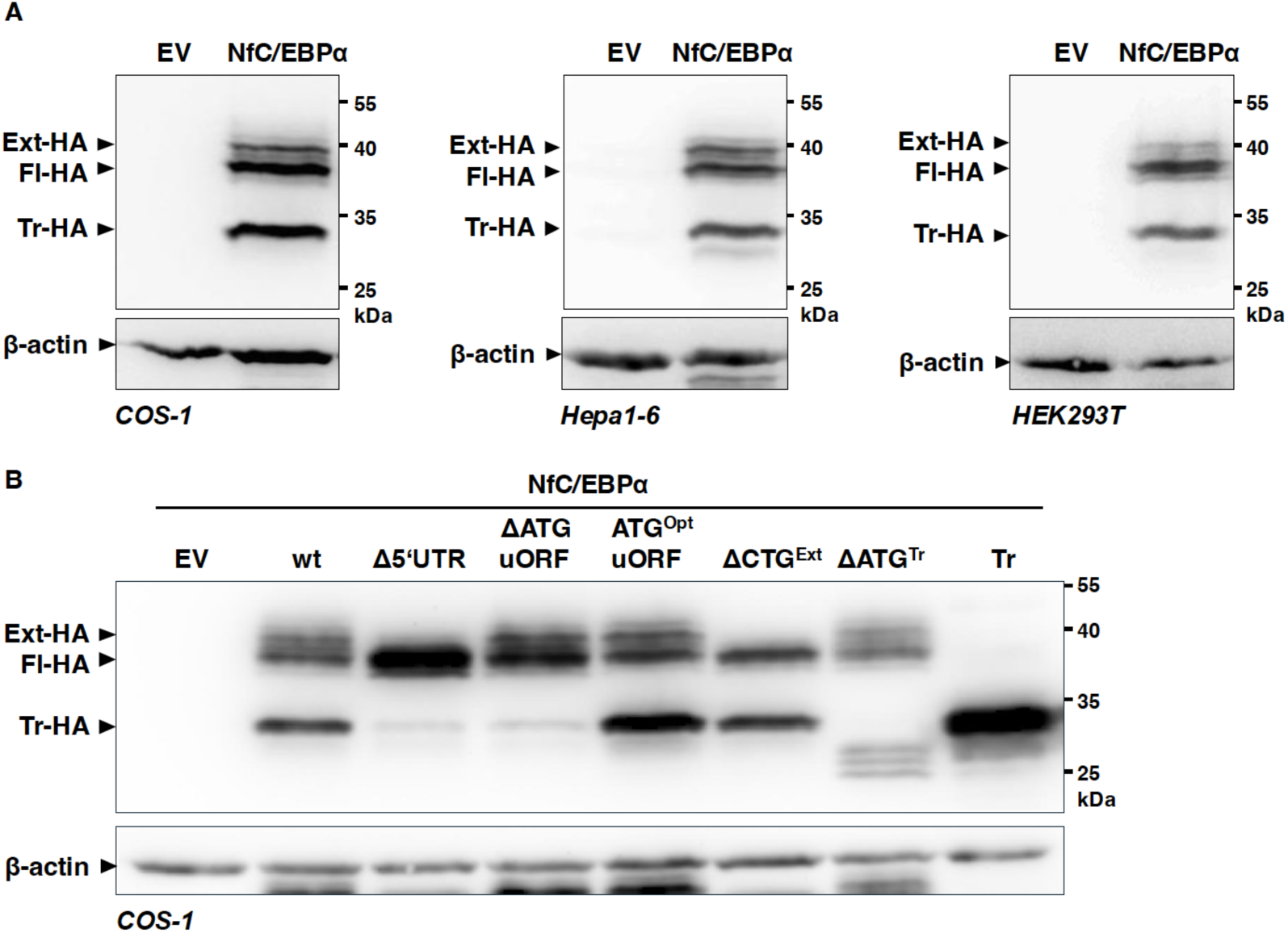
Differential translation of CEBPA mRNA into three NfC/EBPα protein isoforms. (**A**) COS-1, Hepa1-6 and HEK293 cells were transfected with the wild-type (wt) NfC/EBPα-HA expression vector, and protein expression was determined by immunoblotting using anti-HA-antibodies. The NfC/EBPα-HA protein isoforms are labeled as extended (Ext-HA), full-length (Fl-HA), and truncated (Tr-HA). β-actin served as a loading control. (**B**) COS-1 cells were transfected with NfC/EBPα-HA expression vectors containing the following mutations: deletion of the entire 5’UTR (Δ5’UTR), deletion of the uORF-ATG (ΔATG), placing the uORF-ATG in a Kozak sequence for optimal translation initiation efficiency (ATG^Opt^), deletion of the predicted extended-CUG initiation codon (ΔCTG^Ext^), deletion of the double AUG predicted as an initiation site for the truncated isoform (ΔATG^Tr^), and an expression vector for truncated (Tr)-NfC/EBPα only. β-actin served as a loading control.

In summary, the regulation of NfC/EBPα protein isoform expression relies on the same structural elements in the NfCEBPA mRNA and follows the same regulatory rules involving a *cis*-regulatory uORF, as demonstrated in human, rat, mouse, and chicken orthologs (Calkhoven et al., 1994; Calkhoven et al., 2000; Muller et al., 2010).

### Removal of the ΔuORF enhances NfC/EBPα transactivation activity

To investigate whether the truncated NfC/EBPα isoform, expressed from the NfCEBPA mRNA, affects the overall transactivation potential of NfC/EBPα proteins, we co-transfected HEK293T or COS-1 cells with a luciferase reporter vector containing two consensus C/EBP binding sites and the following expression vectors: wild-type NfC/EBPα (wt); a mutant with the upstream CTG initiation codon removed (ΔCTG), preventing expression of Ext-NfC/EBPα; a mutant with the uORF removed (ΔATG^uORF^), reducing Tr-NfC/EBPα in favor of Fl-NfC/EBPα; a mutant with an optimized Kozak sequence at the uORF initiation site (ATG^uORFopt^), increasing Tr-NfC/EBPα at the cost of Fl-NfC/EBPα; and an expression vector for Tr-NfC/EBPα only (**Figure 3B and Figure 4B**). The ΔCTG mutation did not alter the transactivation potential compared to the wild-type NfC/EBPα in both cell lines (**Figure 4A and C**), likely due to the distinct function of Ext-NfC/EBPα in Pol I-mediated rDNA transcription in the nucleolus (Muller et al., 2010). The ΔATG^uORF^ mutation resulted in a significant increase in NfC/EBPα transactivation potential compared to wild-type NfC/EBPα, likely due to diminished expression of the truncated isoform, which normally competes with the extended-and full-length isoforms for DNA binding. In contrast, the ATG^uORFopt^ mutant showed decreased transactivation potential, reflecting the increased expression of the inhibitory truncated isoform. Finally, expression of the Tr-NfC/EBPα alone was unable to activate the transcription reporter above the basic empty vector level (EV) (**Figure 4A and C**). Thus, as observed in other vertebrates, the loss of uORF-mediated Tr-NfC/EBPα isoform expression leads to enhanced NfC/EBPα activity.

**Figure 4.**
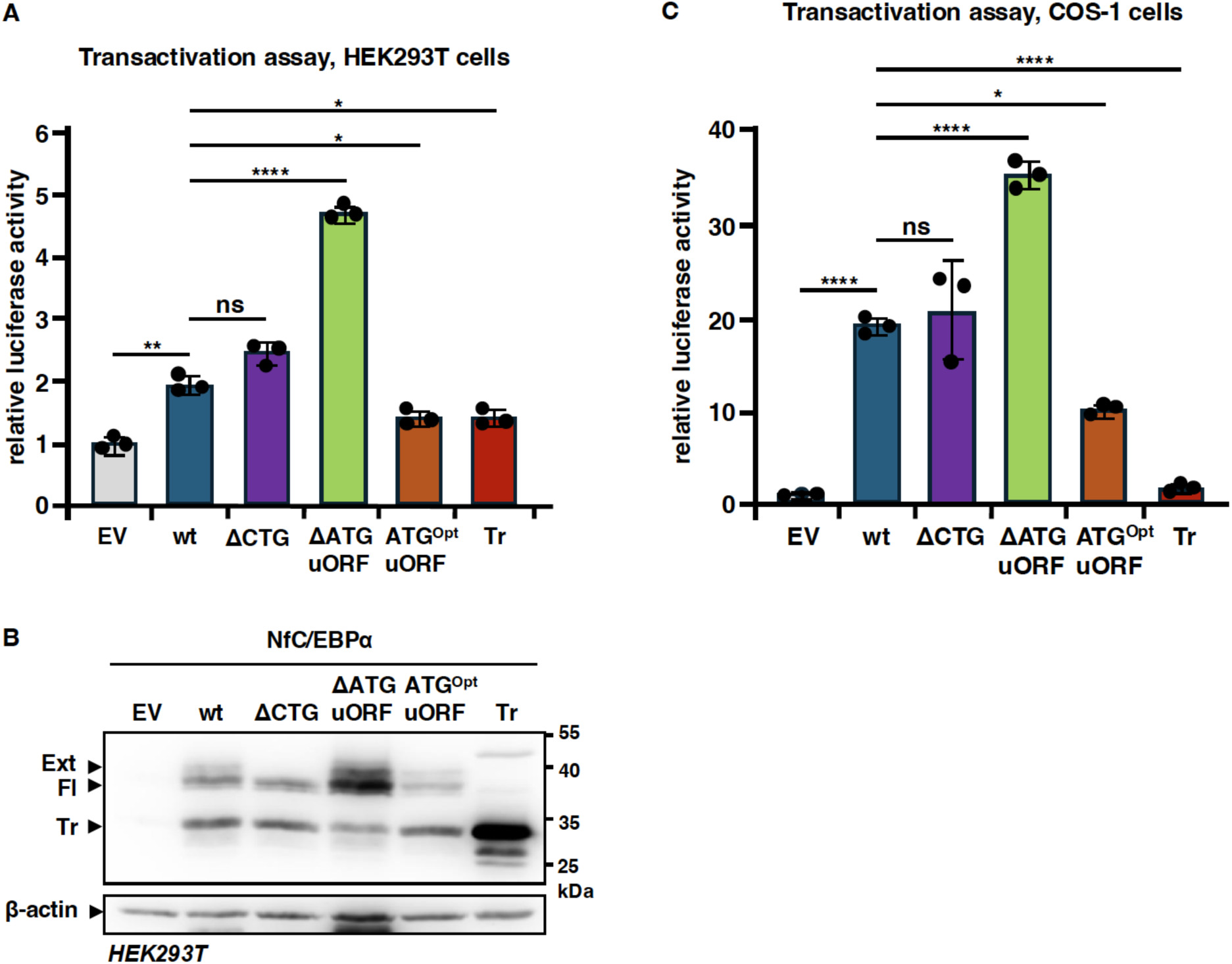
Impact of the CEBPA-ΔuORF mutation on NfC/EBPα transactivation activity. (**A**) Bar graphs showing the fold induction of the 2xC/EBP-binding sites luciferase reporter in HEK293T cells with coexpression of various NfC/EBPα constructs: wild-type (wt), ΔCTG (CTG for Ext-NfC/EBPα removed), ΔATG-uORF (removal of the uORF), ATG^Opt^-uORF (optimizing uORF function), and Tr-NfC/EBPα only. (**B**) Immunoblot analysis of NfC/EBPα construct expression in HEK293T cells, with β-actin serving as a loading control. (**C**) Bar graphs showing the fold induction of the 2xC/EBP-binding sites luciferase reporter in COS-1 cells with coexpression of NfC/EBPα wild-type and mutants. Expression of these constructs in COS-1 cells is shown in Figure 3B. Statistical differences were determined by one-way ANOVA with multiple comparisons. Error bars represent ± SD. *: p<0.05, **: p<0.01, ****: p<0.001.

### Enhanced NfC/EBPα activity extends lifespan in male *N. furzeri*

In the *N. furzeri* strain ZMZ1001, we used CRISPR/Cas9 genome editing to disrupt uORF functionality by mutating uORF-ATG to TTG, creating the *NfCEBPA^ΔuORF^* strain (**Figure 5A**). Successful mutation was confirmed by sequencing (**Figure 5B**) and PCR analysis (**Figure 5C**). Heterozygous males and females were bred to generate homozygous offspring.

**Figure 5.**
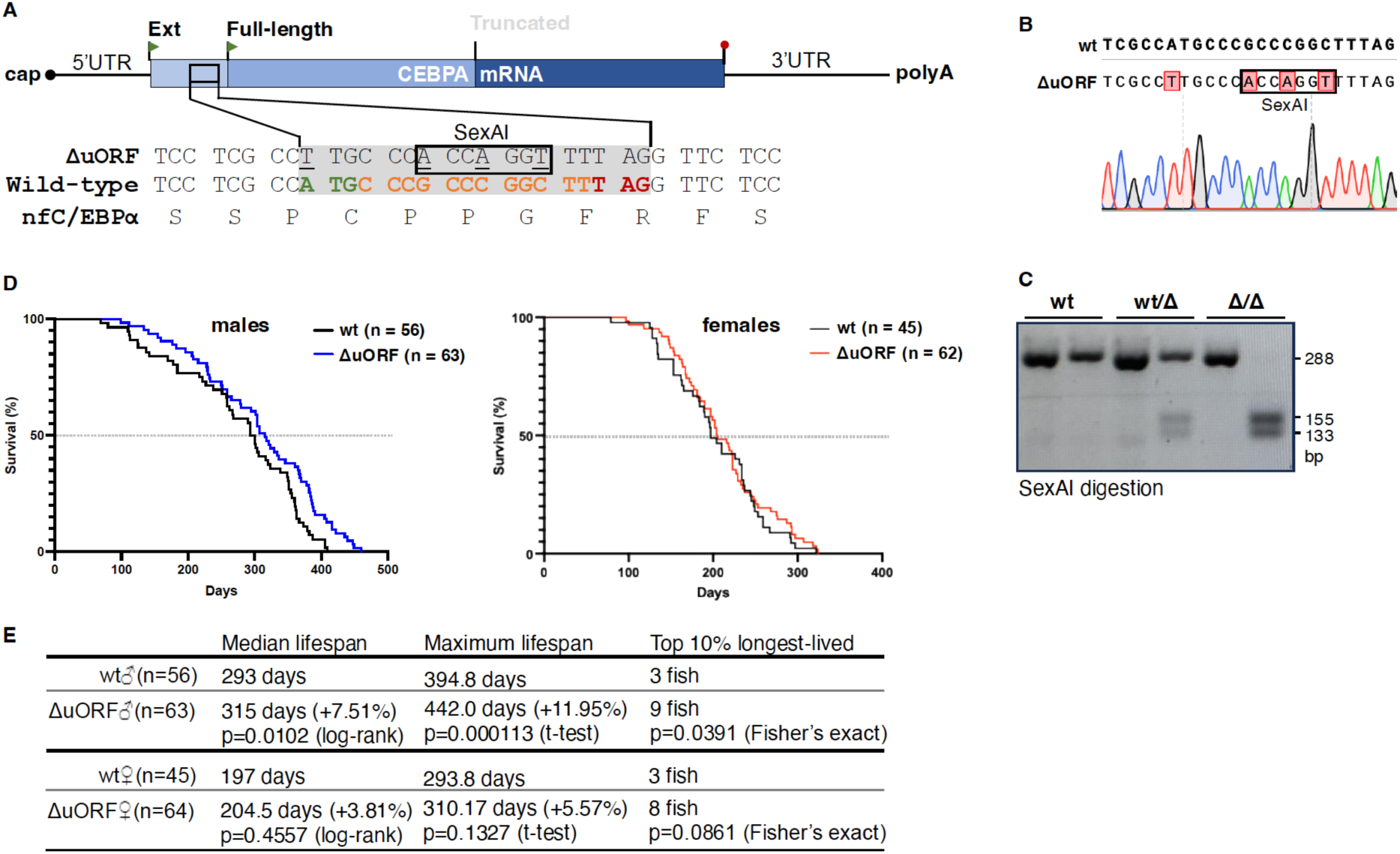
CEBPA-ΔuORF mutation extends the lifespan of male killifish. (**A**) Schematic representation of NfCEBPA mRNA with corresponding cDNA sequences showing mutations that ablate the uORF and introduce a SexAI restriction site for genotyping purposes. Wild-type NfC/EBPα sequences show the +2 out-of-frame uORF sequence highlighted in color. The NfC/EBPα amino acid sequence shown below reveals that the Ext-NfC/EBPα sequence is not affected by the mutation. (**B**) Sequencing result of genomic DNA from ΔuORF homozygous mutant fish. (**C**) PCR analysis of *NfCEBPA* genomic DNA from the wild-type, heterozygous, and homozygous ΔuORF mutants. (**D**) Survival curves of lifespan experiments for wild-type (wt) or *NfCEBPA^ΔuORF^* fish, separated by sex. (**E**) Table summarizing the median lifespan, maximum lifespan, and top 10% longest-lived fish, with statistical analysis as indicated.

To examine lifespan differences between wild-type fish and *NfCEBPA^ΔuORF^* mutants, we monitored offspring from the same heterozygous mating pairs across six female and seven male cohorts. In total, 45 female wt, 62 female ΔuORF, 56 male wt, and 63 male ΔuORF were observed from an age of eight to ten weeks until natural death or termination based on humane endpoint criteria. Analysis of the combined male cohorts revealed a significantly extended lifespan for *NfCEBPA^ΔuORF^* males compared to wild-type males (**Figure 5D and E**). Specifically, the median lifespan increased by 7.5%, whereas the maximum lifespan, defined as the mean lifespan of the top 10% longest-lived fish in each cohort, was extended by nearly 12%. Among the top 10% of longest-lived male fish, nine carried the ΔuORF mutation, compared to only three wild-type fish. No significant difference in lifespan was observed between the female wild-type and *NfCEBPA^ΔuORF^* fish (**Figure 5D and E**). Finally, comparing wild-type killifish, females had shorter lifespans than males, with median lifespans of 197 and 293 days for females and males, respectively (**Figure 5E**). These findings suggest that the ΔuORF mutation, which increases NfC/EBPα transactivation potential, confers a lifespan advantage to male fish of the *N. furzeri* ZMZ1001 strain, particularly during later stages of life, whereas females do not show a similar benefit.

### Enhanced NfC/EBPα activity partially improves healthspan in *N. furzeri*

To assess the healthspan in the lifespan cohorts, we conducted twice-daily visual inspections of each fish, documenting changes in appearance and behavior. **Tables 1A and B** provide an overview of the outcomes, including spontaneous mortality, humane endpoint-based terminations, aging-related phenotypes, and swimming abnormalities, some of which are discussed in further detail below.

**Table 1A.** Mortality, humane endpoint termination, and aging phenotypes.

| sex | genotype | no. | found dead<br>% (no.) | terminated<br>% (no.) | weight loss<br>% (no.)* | dropsy<br>% (no.)* | rectal prolapse<br>% (no.)* | color loss<br>% (no.)<sup>§</sup> | tumors<br>% (no.)<sup>§</sup> |
| --- | --- | --- | --- | --- | --- | --- | --- | --- | --- |
| male | wild-type | 56 | 28.6 (16) | 71.4 (40) | 12.5 (7) | 25.0 (14) | - | 37.5 (21) | 10.7 (6) |
| | $\Delta uORF$ | 63 | 28.6 (18) | 71.4 (45) | 20.6 (13) | 28.6 (18) | - | 33.3 (21) | 4.8 (3) |
| female | wild-type | 45 | 64.4 (29) | 35.6 (16) | 2.2 (1) | 62.5 (11) | 8.9 (4) | 13.3 (6) | 17.8 (8) |
| | $\Delta uORF$ | 62 | 59.7 (37) | 40.3 (25) | 4.8 (3) | 64.0 (16) | 6.5 (4) | 17.7 (11) | 8.1 (5) |

**Table 1B.** Abnormal swimming phenoptypes.

| sex | genotype | no. | belly sliding<br>% (no.)<sup>#§</sup> | floating<br>% (no.)<sup>§</sup> | upside down<br>% (no.)<sup>§</sup> | unbalanced<br>% (no.)<sup>§</sup> | sideways<br>% (no.)<sup>§</sup> | total<br>% (no.)<sup>§</sup> |
| --- | --- | --- | --- | --- | --- | --- | --- | --- |
| male | wild-type | 56 | 21.4 (12) | 5.4 (3) | 10.7 (6) | 3.7 (2) | 16.1 (9) | 57.2 (32) |
| | $\Delta uORF$ | 63 | 20.6 (13) | 4.8 (3) | 3.2 (2) | 1.6 (1) | 11.1 (7) | 41.3 (26) |
| female | wild-type | 45 | - | 6.7 (3) | 2.2 (1) | 6.7 (3) | 8.9 (4) | 24.4 (11) |
| | $\Delta uORF$ | 62 | 3.2 (2) | - | - | - | 12.9 (8) | 16.1 (10) |

\* observed near or at time of humane endpoint termination
<sup>§</sup> observed throughout life
<sup>#</sup> persistent belly sliding for a duration of 7 days or more

Among the spontaneously deceased killifish, approximately one-third were males and two-thirds were females. In contrast, two-thirds of the males were euthanized based on humane endpoint criteria, compared to only one-third of the females (**Table 1A**). The humane endpoint indicators included severe deterioration in general appearance, indicating imminent death, such as abnormal swimming positions and poor body condition, severe abdominal distension (dropsy) with raised scales, visible tumors with progressive growth, often accompanied by poor body condition, markedly reduced movement, and lack of food intake. At the end of life, males exhibited severe weight loss more frequently than females: 12.50% in wild-type and 20.64% in *NfCEBPA^ΔuORF^* males compared to 2.22% in wild-type females and 4.84% in *NfCEBPA^ΔuORF^* females. These data suggest that the *NfCEBPA^ΔuORF^* mutation promotes weight loss in aging fish. The dropsy phenotype was more common in females than in males, with no significant differences between the genotypes. Rectal collapse was observed only in females, and was slightly more frequent in wild-type fish than in *NfCEBPA^ΔuORF^* fish. The incidence of visible tumors was significantly reduced in *NfCEBPA^ΔuORF^* fish, with 10.71% of wild-type males and 17.78% of wild-type females affected compared to 4.76% and 8.07% in *NfCEBPA^ΔuORF^* males and females, respectively. Male killifish, which are naturally more colorful than females, showed age-related loss of color. Although a similar percentage of males showed color loss, this was delayed in the *NfCEBPA^ΔuORF^* males (**Figure 6**).

**Figure 6.**
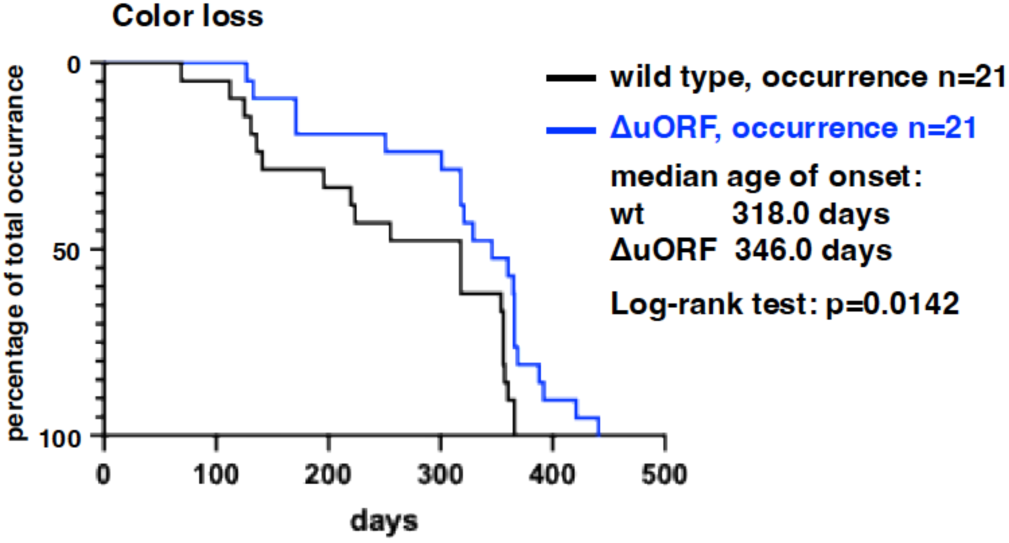
delayed color loss in *NfCEBPA^ΔuORF^* male fish. The curves represent a progressive increase in the percentage of male fish exhibiting color loss throughout their lifespan.

Fish can develop swim bladder disfunctions due to infections, environmental causes, and aging, which manifest as abnormal swimming behaviors. **Table 1B** categorizes five distinct phenotypes: belly sliding, where fish struggle to lift from the bottom of the tank, a condition that, although sometimes observed in young fish and typically resolves within three days, becomes more persistent (over seven days) in older fish; floating, where fish remain at the water surface, often in abnormal body positions; upside down, with the belly facing upwards; unbalanced, meaning that the position of the head is either lower or higher than the tail; and sideways, when fish swim tilted to one side at various angles. Overall, the frequency of abnormal swimming behaviors was significantly higher in males than in females (**Table 1B**). Additionally, the incidence of these behaviors was lower in *NfCEBPA^ΔuORF^* males and slightly lower in *NfCEBPA^ΔuORF^* females, compared to wild-type fish (**Table 1B**). In *NfCEBPA^ΔuORF^* males, the occurrence of abnormal swimming behaviors was significantly delayed compared to wild-type males (**Figure 7A**), which was also observed for belly sliding alone as the largest contributor to abnormal swimming (**Figure 7B**). No significant differences in timing were found among the female fish (**Figure 7C**). Together, these results suggest that while swim bladder function is more susceptible to age-related decline in male fish, this decline is delayed in *NfCEBPA^ΔuORF^* males, indicating a potential protective effect of the ΔuORF mutation on swim bladder function as fish age.

**Figure 7.**
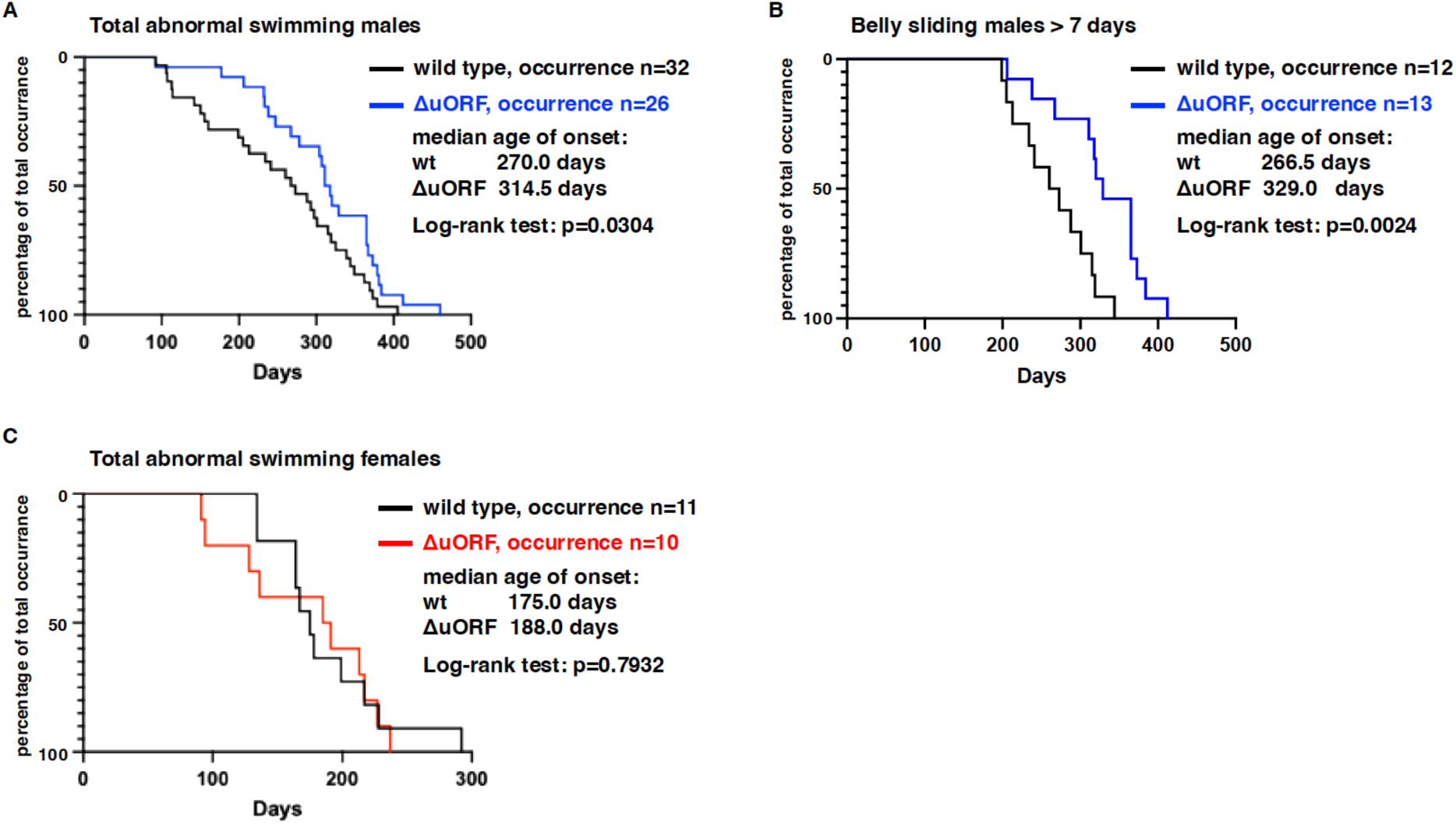
CEBPA-ΔuORF mutation delays abnormal swimming behavior in males. The curves represent the progressive increase in the percentage of fish exhibiting (**A**) total abnormal swim behaviors in males, (**B**) belly-sliding in males, and (**C**) total abnormal swimming in females.

## DISCUSSION

Here, we demonstrate that translation of the *NfCEBPA* gene transcript in the African turquoise killifish *Nothobranchius furzeri* produces three distinct protein isoforms: extended (Ext)-, full-length (Fl)-, and truncated (Tr)-NfC/EBPα. By identifying the translation initiation codons involved, we established that Tr-NfC/EBPα expression depends on the integrity of a small upstream open reading frame (uORF). These features mirror those reported for *CEBPA* orthologs in mice, rats, chickens, and humans, underscoring the evolutionary conservation of the *CEBPA* genes and their C/EBPα protein products across a wide range of vertebrates. Disruption of the *NfCEBPA* uORF in killifish by CRISP/Cas9 mediated genomic modification significantly extended the median lifespan of the male *NfCEBPA^ΔuORF^* fish by +7.51% and the maximum lifespan by +11.95%, along with delays and reductions in various age-related phenotypes. Notably, female *NfCEBPA^ΔuORF^* killifish did not show similar improvements. Despite testing multiple anti-C/EBPα antibodies targeting epitopes that partially overlap with the NfC/EBPα sequences, none demonstrated sufficient reactivity against endogenous NfC/EBPα. However, numerous experiments with C/EBPα and C/EBPβ of other species have consistently demonstrated that uORF-dependent regulation operates identically in both cell culture and whole organisms.

The sex-specific effects on health and lifespan observed in *NfCEBPA^ΔuORF^* killifish remind of the findings in mice carrying a mutation in the uORF of the related *CEBPB* gene (*CEBPB^ΔuORF^* mice). From CEBPB mRNA, three protein isoforms are translated into C/EBPβ-LAP1/2 and the N-terminally truncated isoform C/EBPβ-LIP (Calkhoven et al., 2000; Wethmar, Begay, et al., 2010). Disruption of the *CEBPB* uORF in mice enhances C/EBPβ transactivating activity, resulting in improved metabolic profiles similar to those observed under calorie restriction. In addition, the mutation helps preserve motor coordination and immune functions in older *CEBPB^ΔuORF^* mice compared to their wild-type littermates in both sexes. However, an extension of the median lifespan of 20.6% and a maximum lifespan of 10% was only observed in females (Muller et al., 2018; Muller et al., 2022; Zidek et al., 2015). Additionally, impaired demethylation of C/EBPβ-regulated super-enhancers, which prevents C/EBPβ binding, has been linked to premature aging in mice (Schafer et al., 2018). Together with the data presented here, these findings underscore the essential role of both C/EBPα and C/EBPβ in maintaining health and resilience throughout aging (Niehrs & Calkhoven, 2020).

Within the C/EBP transcription factor family, only *CEBPA* and *CEBPB* share the unique features of being intronless, containing a cis-regulatory uORF, and utilizing three alternative translation initiation sites to produce three distinct protein isoforms of C/EBPα and C/EBPβ (Ramji & Foka, 2002; Wethmar, Smink, & Leutz, 2010). Comparative analysis of mRNA sequences across vertebrates has revealed remarkable conservation in the distribution, position, and relative strength of these initiation sites, highlighting their evolutionary importance. Studies have shown that uORF-driven expression of truncated-C/EBPα (p30) and C/EBPβ-LIP are tightly regulated by the mTORC1-4E-BP and eIF2α kinase pathways (Calkhoven et al., 2000; Zidek et al., 2015), linking these C/EBPs to cellular nutrient and stress signaling. Either inhibition of mTORC1 signaling or activation of eIF2α kinases results in suppression of truncated C/EBPα (p30) and C/EBPβ-LIP expression. These shared regulatory mechanisms suggest that both C/EBPα and C/EBPβ act as integrators of nutrient, growth factor and stress signals to orchestrate a transcriptional response (Niehrs & Calkhoven, 2020). Furthermore, Fl-C/EBPα function is connected to Acetyl-CoA and NAD^+^ metabolism through its acetylation by the acetyltransferase p300 and deacetylation by SIRT1 (Zaini et al., 2018). In response to glucose deprivation, increased NAD+ levels activate SIRT1, which deacetylates C/EBPα. Hypoacetylated C/EBPα shifts the downstream transcriptional response towards genes involved in mitochondrial biogenesis and respiration.

Notably, the most prominent phenotype in the *NfCEBPA^ΔuORF^* fish observed here was a reduced incidence of visible cancers, suggesting a similar prominent role of C/EBPα in cancer development, as has been observed in *CEBPB^ΔuORF^* mice. C/EBPβ-LIP deficiency reduces tumor incidence (Muller et al., 2018), whereas its overexpression increases tumor incidence in mice (Begay et al., 2015). Furthermore, C/EBPβ-LIP is highly expressed in several human cancers, including breast cancer, ovarian cancer, colorectal cancer, and anaplastic large cell lymphoma (Jundt et al., 2005; Quintanilla-Martinez et al., 2006; Rask et al., 2000; Sundfeldt et al., 1999; Zahnow, Younes, Laucirica, & Rosen, 1997), and C/EBPβ-LIP overexpression induces cellular transformation *in vitro* and *in vivo* (Calkhoven et al., 2000; Zahnow et al., 1997). Mechanistically, C/EBPβ-LIP supports oncogenesis through various pathways. For instance, it induces a shift towards cancer-specific metabolic reprogramming by enhancing glycolysis and mitochondrial respiration, mediated in part by the regulation of the let-7/LIN28B circuit (Ackermann et al., 2019). Additionally, C/EBPβ-LIP promotes the activity of the malate-aspartate shuttle, ensuring NADH/NAD^+^ homeostasis in cancer cells (Ackermann et al., 2023). Beyond metabolic reprogramming, C/EBPβ-LIP has been shown to stimulate cancer cell migration and invasion in cell culture, suggesting a crucial role in metastasis (Matherne, Phillips, Embrey, Burke, & Machado, 2023; Sterken et al., 2022).

The truncated isoform of C/EBPα (C/EBPα-p30) has also been implicated in cancer development. In acute myeloid leukemia (AML), mutations in the *CEBPA* gene frequently generate premature stop codons, preventing the expression of the extended (Ext-C/EBPα) and full-length (FL-C/EBPα/ C/EBPα-p42) isoforms, while leaving the C/EBPα-p30 isoform unaffected (Nerlov, 2004). Experimental studies in mice demonstrated that replacing the *CEBPA* allele with sequences exclusively expressing C/EBPα-p30 induces AML with full penetrance, suggesting that C/EBPα-p30 acts as an oncogene (Bereshchenko et al., 2009; Kirstetter et al., 2008). Furthermore, overexpression of C/EBPα-p30 in 3T3-L1 adipoblasts induces cellular transformation (Calkhoven et al., 2000). Mechanistically, C/EBPα-p42 exerts strong anti-proliferative activity, which is dominant over the loss of tumor suppressors, such as p53 or Rb (Hendricks-Taylor & Darlington, 1995; Muller et al., 1999). In contrast, the truncated isoform lacks this tumor-suppressive function, and by competing with C/EBPα-p42 for binding to target gene promoters and enhancers it facilitates cell proliferation. The specific downstream mechanisms and physiological processes by which C/EBPα induces cell proliferation and contributes to cancer development require further investigation.

Both male and female *NfCEBPA^ΔuORF^* fish showed a notable reduction in abnormal swimming behavior compared to their wild-type counterparts. These phenotypes, which are often linked to swim bladder dysfunction, can arise from infections, environmental influences, or age-related decline. The reduced incidence of such behaviors in mutant fish suggests an enhanced resilience to these challenges. Although this finding is limited in scope, it aligns with observations in *NfCEBPB^ΔuORF^* mice, which display increased resistance to ulcerative dermatitis, a condition prevalent in C57BL/6 mice (Muller et al., 2018). Additionally, older *NfCEBPB^ΔuORF^* mice display a remarkably preserved motor coordination and a more youthful immune profile, indicated by increased memory/naïve T-cell ratios, highlighting the broader health benefits conferred by uORF disruption in these transcription factors.

In this study using killifish and in studies using mice, sex-specific phenotypes in response to genetic modifications were observed. Such differences in responses to mutations or treatments are often poorly understood. In laboratory animal models, it is important to consider that genetic background plays a significant role. For example, the genetic makeup of the *N. furzeri* ZMZ1001 strain used in this study may substantially contribute to the observed sex-specific responses. Similarly, studies in genetically diverse mice under various calorie restriction regimens have revealed a strong dependence on genetic background for phenotypic variation. In addition, hormonal interactions, both systemic and at the level of nuclear hormone receptor-C/EBP interactions, likely play essential roles in mediating sex-specific effects.

This study aimed to confirm the evolutionary conservation of C/EBPα regulation, focusing on its primary mRNA structure, while also providing novel insights into how altering NfC/EBPα protein isoform expression impacts health and lifespan. These findings, along with research on C/EBPβ in mice, underscore the importance of further investigation into the role of C/EBP transcription factors in promoting resilience against age-related conditions and diseases.

The parallels between C/EBPα and C/EBPβ suggest shared and distinct roles in the regulation of cellular processes that contribute to aging and longevity. Key unanswered questions remain, including identifying the specific biological processes uniquely regulated by C/EBPα or C/EBPβ, and those where their functions overlap. Investigating how these transcription factors integrate nutrient and stress signaling pathways, such as those involving mTORC1, eIF2α-kinases, and sirtuins, could reveal how environmental and metabolic cues influence their activity. Additionally, the interplay between C/EBPs and other transcriptional regulators, such as nuclear hormone receptors, deserves more attention to understand their roles in cellular homeostasis.

Although it is challenging to pharmacologically target transcription factors because of their structural properties, the signaling pathways that regulate their activity and/or expression represent promising therapeutic avenues (Zaini et al., 2017). Identifying upstream modulators of C/EBP function and dissecting their mechanistic roles is essential for developing strategies to enhance resilience against aging and age-related diseases. Finally, expanding research in diverse model organisms and human systems is also critical for translating these findings into clinical applications.

## MATERIALS AND METHODS

### DNA constructs

The *CEBPA* cDNA from *N. furzeri* ZMZ1001 strain corresponding to the 5’UTR and the whole coding region (wt-C/EBPα), and the Δ5’UTR construct lacking the 5’UTR sequence were amplified using genomic DNA as a template, and fused at their C-terminus to the HA epitope sequence and cloned into the pCDNA3.1 expression vector (Invitrogen) by Gibson Assembly using an NEBuilder HiFi DNA Assembly kit from NEB with the primers listed in the table below. The Fl-, Tr-, ΔATG-uORF-, ATG^Opt^-uORF-and ΔATG^Tr^-C/EBPα pCDNA3.1 constructs were generated using whole plasmid PCR amplification, phosphorylation of the PCR fragments with Polynucleotidkinase, and blunt ligation (re-circulation) (see table below for primers). The ΔCTG^Ext^-C/EBPα pCDNA3.1 construct was generated by exchanging the N-terminus of the C/EBPα-wt pCDNA3.1 construct with a mutated fragment obtained by PCR amplification, using NheI and AgeI restriction endonucleases. To subclone NfC/EBPα expression constructs into the pSG5 expression vector, C/EBPα-wt and mutant sequences were amplified by PCR using the corresponding pCDNA3.1 based constructs and cloned into the EcoRI site of the pSG5 vector (Stratagene). The sequence of all constructs was confirmed by Sanger sequencing.

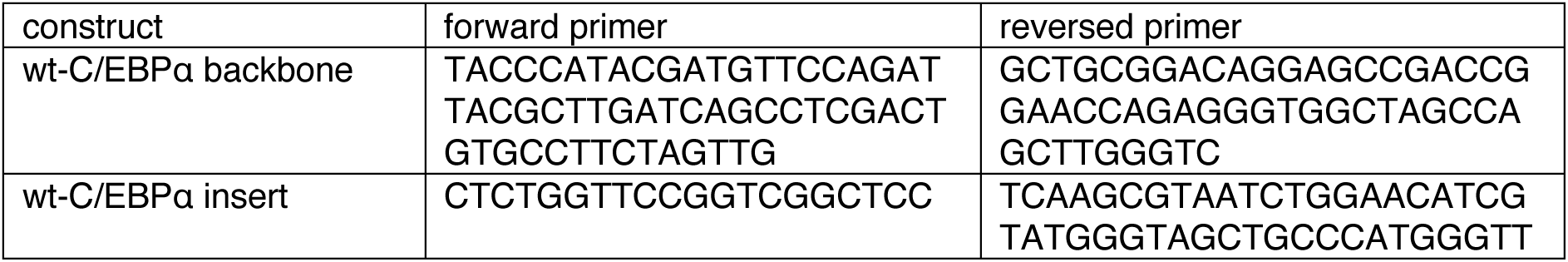
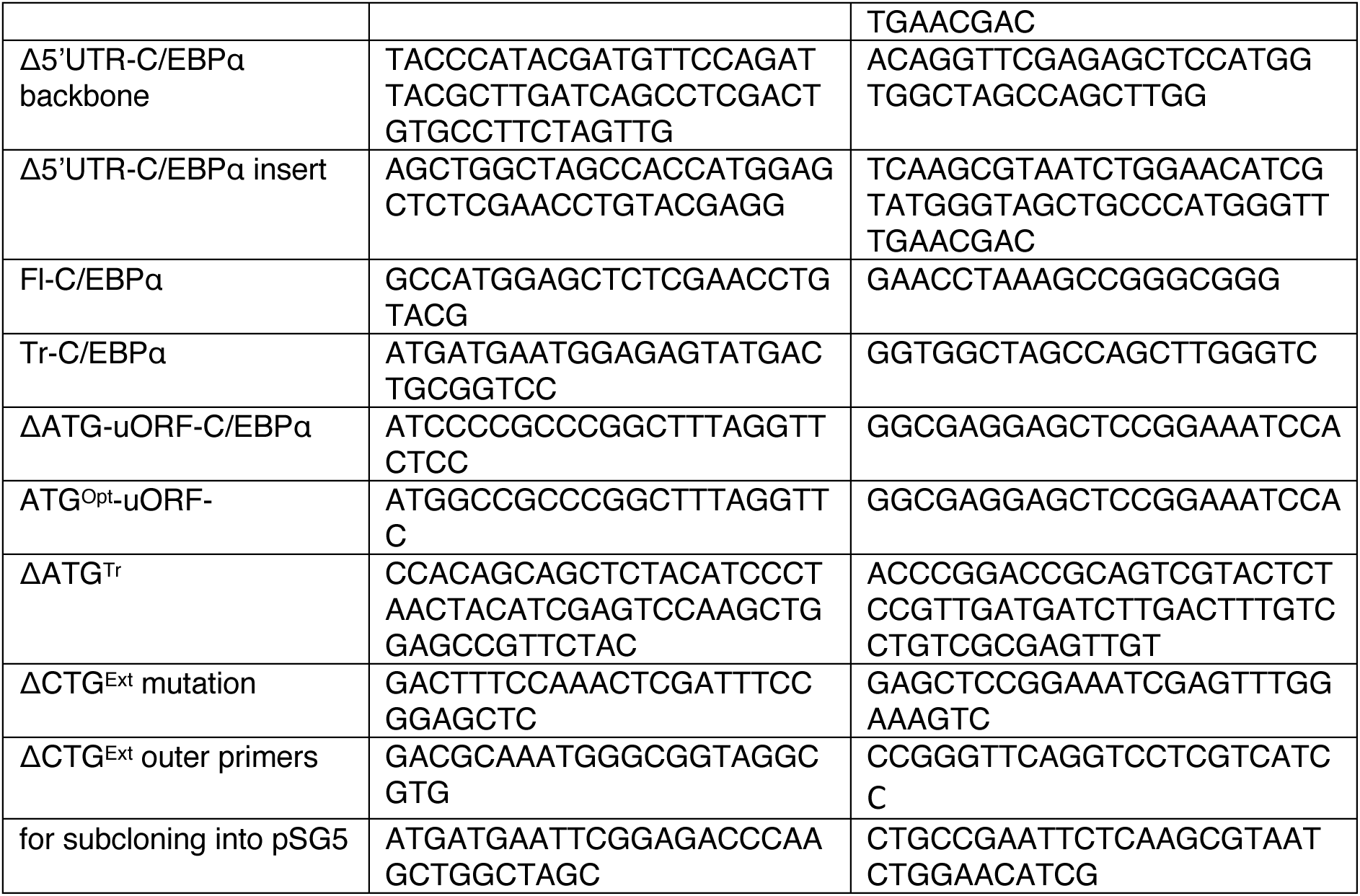

### Cells

COS-1 cells were cultured in DMEM/F12 medium supplemented with 5% FCS, 10mM HEPES, and penicillin/streptomycin. HEK293T cells and Hepa1-6 cells were cultured in DMEM medium supplemented with 10% FCS, 10mM HEPES, and penicillin/streptomycin. All cell lines were maintained at 37°C in a humidified incubator with 5% CO_2_.

### Transfection

COS-1 cells were transfected with 5 μg of pSG5-based expression constructs using the DEAE-dextran/chloroquine method described in (Gonzalez & Joly, 1995). HEK293T cells were transfected with 2.5 μg of pCDNA3.1-based expression constructs using the calcium phosphate precipitation method. Hepa1-6 cells were transfected with 4 μg of pCDNA3.1-based expression constructs using 12 μl Fugene HD (Promega), following the manufacturer’s protocol.

### Luciferase reporter assay

Cells were seeded into a 96-well plate at a density of 5000 cells (HEK293T) or 6000 cells (COS-1) per well. After 24 hours, the cells were transfected with 600 ng of pCDNA3.1-based (HEK293T) or pSG5- based (COS-1) expression vectors, 300 ng of a firefly luciferase reporter vector containing two consensus C/EBP binding sites (pM82; lacking the AP-1 binding site (Sterneck, Muller, Katz, & Leutz, 1992)), 100 ng of pLG4 renilla luciferase vector for normalization, and 2.5 μl (HEK293T) or 3 μl (COS-1) Fugene HD transfection reagent (Pomega). The transfection mix was divided over three cell containing wells for triplicate values. After 48 hours, luciferase activity was measured using Dual-Glo luciferase assay system (Promega) according to the manufacturer’s instructions. Luminescence was detected with a GloMax-Multi detection system (Promega).

### Immunoblotting

Cells were washed twice with cold PBS and harvested with a cell scraper. Cells were lysed in 50 mM Tris pH 8.0, 150 mM NaCl, 0,5% sodium deoxycholate, 0.1% SDS and 1% Triton-X 100, supplemented with protease and phosphatase inhibitors (Roche), and sonicated. Equal amounts of protein were separated by SDS-PAGE and transferred to 0.2 μm PVDF membranes by a Trans-Blot Turbo System (Bio-Rad). Detection was performed by using the following antibodies: anti-HA (MMS-101R; Covance, 1:1000), anti-β-Actin (clone C4, 691001; MP Biomedicals, 1:10000), HRP-conjugated anti-mouse (GE Healthcare, 1:5000), and the lightning Plus ECL reagent (Perkin Elmer). For reprobing, membranes were incubated with restore western blot stripping buffer (Thermo Fisher). The detection was performed with the Image Quant LAS 800 Imager (GE Healthcare).

### Generation of *NfCEBPA^ΔuORF^* fish

Chemically modified guide crRNAs were synthesized by Integrated DNA Technologies (IDT). The crRNAs were designed using the IDT alt-R guide RNA design tool, and three overlapping guide RNAs were selected (https://eu.idtdna.com/site/order/designtool/index/CRISPR_CUSTOM). A 105 bp oligonucleotide with 54- and 38-bp homology arms, respectively, and two consecutive phosphorothioate modifications on both the 5’ and 3’ ends was synthesized by IDT to serve as the repair template. The target regions of the guide crRNAs and the repair template are listed in the table below.

Injection mixes were prepared as per protocol outlined at the IDT web site (https://sfvideo.blob.core.windows.net/sitefinity/docs/default-source/user-submitted-method/crispr-cas9-rnp-delivery-zebrafish-embryos-j-essnerc46b5a1532796e2eaa53ff00001c1b3c.pdf?sfvrsn=52123407_10). Briefly, crRNA and tracrRNA were diluted to 3 μM (1 μM for each crRNA) in IDT Nuclease-Free duplex buffer and annealed by heating at 95 °C for 5 min, followed by cooling to room temperature. The solution was mixed with an equal volume of 0,5 μg/μl Cas9 protein (alt-R, IDT) in Cas9 working buffer (20 mM HEPES, 150 mM KCl, pH 7.5). Injections were performed in one-cell stage eggs as described in (Valenzano, Sharp, & Brunet, 2011). A total of 697 eggs were injected, with 100 animals surviving to adulthood. These were in-crossed, and eggs from each breeding pair were collected and genotyped as described below. Three founders with the correctly edited base pairs were identified. The combined offspring of the 39 F1 animals were genotyped using tissues obtained by fin clipping. Four heterozygoes males were identified and subsequently outcrossed twice.

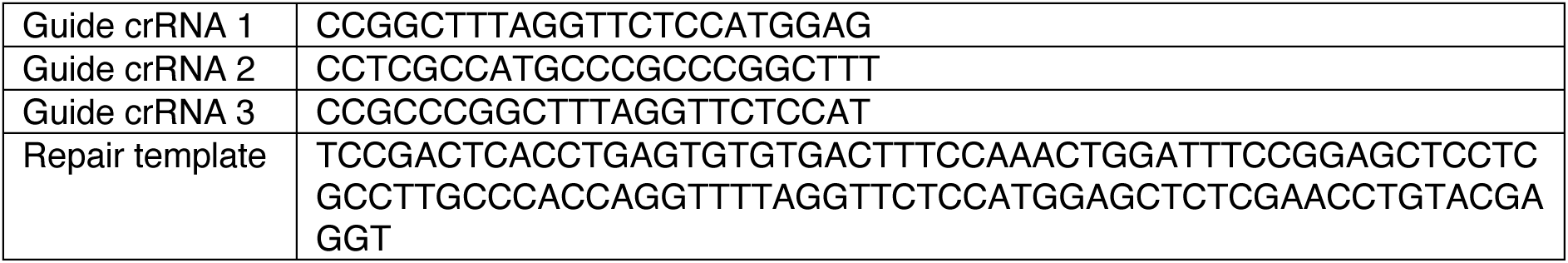

### Genotyping

Sampled material (eggs or fin clips) was heated in 50 μl 0.05 M NaOH at 95°C for 10 minutes. The reaction was neutralized by adding 5 μl 1 M Tris-HCl (pH 8.0). The 289 bp region of interest was amplified using the primers: forward, 5’-CTC TTC GTT CCA ACA CAA AGT G-3’ and reverse, 5’-GGT CTA TGG AGG TCT CGT TGT C-3’. For restriction analysis, half of the PCR product was digested with the restriction enzyme SexAI. Both the untreated and digested samples were then subjected to DNA electrophoresis on adjacent lanes for analysis.

### Fish maintenance

Fish of the *Nothobranchius furzeri* strain ZMZ-1001 were maintained at 28°C on a 12-hour light/12-hour dark cycle in 3.5-liter tanks that were connected to a recirculation water system. The fish were fed with frozen bloodworms (Chironomus, purchased from Dutch Select Food) twice a day during the week and once a day during the weekend. At six to seven weeks of age, fin clips were taken for genotyping. After genotyping, female fish were combined to obtain four to five fish with the same genotype in one tank, while male fish stayed separated. For breeding, single heterozygous males and one to three heterozygous females between 8 and 10 weeks of age were placed together in a 3.5-liter tank containing a tray with autoclaved fine sand for egg collection. Eggs were collected from the sand trays by sieving and transferred to a petri dish containing filter-sterilized tank water. Eggs were then bleached by incubation for 10 minutes in sterile tank water containing 0.025% sodium hypochlorite, washed twice with sterile tank water, and transferred to a new Petri dish with sterile tank water containing gentamycin (10μg/ml) and Methylene Blue (1:20000), and incubated at 28°C. 48 hours after embryos reached diapause II, eggs were transferred to Petri dishes containing autoclaved Whatman paper soaked in humic acid (1g/l sterile tank water) and further incubated at 28°C until embryos were fully developed. Hatching was induced by transferring the eggs to cold (4°C) humic acid (1g/l sterile tank water) and incubation at 28°C for 4 hours after which the hatched fish were placed together in tanks (max. 50 fish per tank). Feeding started the next day as described above. Two weeks after hatching, fish were transferred to individual tanks.

### Lifespan experiment

Wild-type and homozygous *NfCEBPA^ΔuORF^* mutant fish that were obtained from the same breeding pairs using heterozygous parents were included in the experiment after successful genotyping at eight to ten weeks of age. The lifespan of the fish was determined from the day of hatching until natural death occurred or until the fish had to be terminated because humane endpoint criteria were met. In the latter case, fish were terminated by exposure to ice water for five minutes. Dead fish (both naturally deceased and terminated) were frozen at −20°C, and fin clips were taken again for re-genotyping. Difference in survival was determined using Prism software from GraphPad. The experiments were approved by the Central Commission of Animal Experiments of the Dutch State (Centrale Commissie Dierproeven, CCD, license no AVD1050020197765).

### Determination of aging phenotypes

The health and behavioral state of every fish in the lifespan experiment was examined and noted twice a day (once a day on the weekend) by experienced animal caretakers, and additionally twice a week by a scientist involved in the project. The remarks from these examinations provided the basis for the healthspan analysis, in which the age of onset of different health parameters was determined. These health parameters included visible tumors on the outside of the fish; either outgrowth of cell mass with an increase in size or an increasing area of black pigmentation in the fish body, excluding the tail fin. Potential tumors were not verified by pathological examination. In addition, rapid loss of color, rectal prolapse, dropsy, weight loss (based on visible examination), and abnormal swimming phenotypes were monitored. Within the abnormal swimming phenotypes, we discriminated between belly sliding (fish struggle to lift from the bottom of the tank) for a duration of at least seven days, floating (fish remain at the water surface), swimming upside down (belly facing upwards), unbalanced (position of the head is higher or lower than the tail), or sideways (fish swim tilted to one side at various angles).

## AUTHOR CONTRIBUTIONS

*Study design:* C.M, E.B. and C.F.C. *Study conduct and data collection:* C.M., J.S. M., G.K. and J.H. *Data analysis and interpretation:* C.M., J.S.M. and C.F.C. *Manuscript writing:* C.M., E.B. and C.F.C.

## ACKNOWLEDGEMENTS

We would like to express our gratitude to Pieter Stijvers, Ralph Reinders and Jessica M. de Vries of the animal facility (CDP) at the UMCG for their invaluable assistance. From the Leibniz Institute on Aging – Fritz Lipmann Institute (FLI), we thank Dario R. Valenzano for advice in establishing the killifish facility and providing the fish. We appreciate the efforts of Dagmar Kruspe and Christoph Englert from the FLI in attempting to transfect a killifish cell line.

